# An RGB based deep neural network for high fidelity Fusarium head blight phenotyping in wheat

**DOI:** 10.1101/2023.09.20.558703

**Authors:** Julian Cooper, Chuan Du, Zach Beaver, Ming Zheng, Rae Page, Joseph R. Wodarek, Oadi Matny, Tamas Szinyei, Alejandra Quiñones, James A. Anderson, Kevin P. Smith, Ce Yang, Brian J. Steffenson, Cory D. Hirsch

## Abstract

Fusarium head blight (FHB) in wheat is an economically important disease, which can cause yield losses exceeding 50% and the causal pathogen that infects spikes produces harmful mycotoxins. Breeding for host resistance remains the most effective disease control method; but time, labor, and human subjectivity during disease scoring limits selection advancements. In this study we describe an innovative, high-throughput phenotyping rover for capturing in-field RGB images and a deep neural network pipeline for wheat spike detection and FHB disease quantification. The image analysis pipeline successfully detects wheat spikes from images under variable field conditions, segments spikes and diseased tissue in the spikes, and quantifies disease severity as the region of intersection between spike and disease masks. Model inferences on an individual spike and plot basis were compared to human visual disease scoring in the field and on imagery for model evaluation. The precision and throughput of the model surpassed traditional field rating methods. The accuracy of FHB severity assessments of the model was equivalent to human disease annotations of images, however individual spike disease assessment was influenced by field location. The model was able to quantify FHB in images taken with different camera orientations in an unseen year, which demonstrates strong generalizability. This innovative pipeline represents a breakthrough in FHB phenotyping, offering precise and efficient assessment of FHB on both individual spikes and plot aggregates. The model is robust to different conditions and the potential to standardize disease evaluation methods across the community make it a valuable tool for studying and managing this economically significant fungal disease.

## Introduction

Fusarium head blight (FHB) is an economically significant disease of wheat (*Triticum aestivum*), caused by the fungal pathogen *Fusarium graminearum.* Yield losses from FHB can exceed 50% under favorable conditions, and in the U.S. 28 million tons of wheat grain are estimated to be lost annually to this disease (Savary *et al*., 2019; Friskop *et al*., 2021). Additionally, the causal organisms of FHB can produce mycotoxins which are harmful to humans and animals and reduce the market value of the grain (McMullen *et al*., 2012). Fungicides have been utilized to control FHB outbreaks, but their use is limited by the cost of the chemicals and short window surrounding heading when they can be applied (Beyer *et al*., 2006). To help guide fungicide use decisions with risk assessments, local weather-based forecasting models have been developed (Shah *et al*., 2013). Numerous forecasting models have been developed around the world, including the National FHB risk model commonly used throughout North America (Cowger *et al*., 2020), but they are often highly specific to their region of origin which limits widespread use of a single model. Crop rotation and residue control are also forms of cultural control, but regional changes are required to limit infection from nearby inoculum sources (Dill-Macky and Jones, 2000). The effectiveness of chemical and cultural control methods varies and favorable weather conditions can still lead to disease epidemics (Torres *et al*., 2019). Breeding and growing resistant germplasm can consistently reduce FHB infection by 30-50% and is the most effective strategy for both short and long-term management of FHB to protect global food supply and food safety (McMullen *et al*., 2012; Buerstmayr *et al*., 2020).

Plant resistance to FHB is quantitative, and effective in-field phenotyping is necessary to identify the many small effect loci that confer meaningful resistance (Serajazari *et al*., 2023). The symptoms of the disease can often be mistaken for normal plant maturity, which makes phenotyping subjective and increases rater variation. In addition, variable severity between spikes within a plot increases the time and labor necessary for disease ratings to accurately capture host response (Kant *et al*., 2011). Research groups use different rating methods to capture disease variability and host response. By averaging the number of infected kernels from multiple individual spikes or giving a 0-100% disease severity score to a random sample of spikes in a plot, researchers are able to limit subjectivity, but time and labor requirements are high (Huang *et al*., 2018; Serajazari *et al*., 2023). In contrast, assigning a 0-100% aggregate disease severity based on a full plot visual inspection reduces time and labor, but introduces greater human bias to disease scoring (Talas *et al*., 2012). Furthermore, the lack of a standard measurement for FHB severity limits the usefulness of data sharing across the community (Leonelli *et al*., 2017). A standardized FHB disease assessment method that is objective, high-throughput, high resolution, and less labor intensive would increase the selection accuracy for resistant germplasm. It would also increase the scope and scale of disease testing, and fill the need to develop standard phenotyping protocols, nomenclature, annotation, and databases to enable comparative phenomics across this internationally important disease.

Many types of imaging sensors and platforms are used for field-based high-throughput phenotyping (Sweet *et al*., 2022). Satellites, unoccupied aerial vehicles, and rovers have been used with different types of sensors for disease detection (Raza *et al*., 2020; Sandino *et al*., 2018; Su *et al*., 2020). Red-Green-Blue (RGB) cameras capture visible light from 400-700 nm and can be used to detect disease symptoms (Amarasingam *et al*., 2022). Hyperspectral cameras capture wavelengths across the electromagnetic radiation spectrum and have been used to identify plant disease before visible symptoms occur (Behmann *et al*., 2014). Various non-spectral sensors are also commonly employed in agriculture and plant pathology settings, including light detection and ranging (LiDAR) and thermal devices (Eitel *et al*., 2014; Kefauver *et al*., 2017). The choice of platform and sensor to use for phenotyping is often dictated by cost, the features of the trait of interest, and scale at which phenotyping needs to occur. The *Fusarium* species causing FHB infect wheat spikes, which are relatively small plant organs; thus, using high resolution terrestrial phenotyping is thought to be an effective method for screening individual plants compared to aerial or atmospheric imaging. In addition, the large scope and scale of FHB phenotyping needs in research and breeding groups favors inexpensive and easy methods to deploy RGB sensors rather than more expensive and complex options (Adão *et al*., 2017).

Various methods have been used in wheat for high-throughput phenotyping of FHB (Alisaac and Mahlein, 2023). Using different deep learning architectures, spike and disease segmentation has been performed based on RGB images from commercial cameras (Gu *et al*., 2020; Hong *et al*., 2022; Rößle *et al*., 2023). General classification of diseased versus healthy wheat spikes has been performed using multiple types of machine learning models. A classification model based on the AlexNet research network classified preprocessed diseased spike images, taken from controlled in-door settings, into six ordinal classes based on severity. The use of high resolution images taken at 90 degrees to the spike during grain fill produced the best model accuracy (Gu *et al*., 2020). In-field based FHB phenotyping has also been of interest. Using deep learning architecture based on You Only Look Once v4 and MobileNet, FHB detection of wheat spikes was performed on images taken with an RGB camera in a field setting with 94% accuracy (Hong *et al*., 2022). Detecting diseased wheat spikes is important for field scouting but has limited applications when studying host–pathogen interactions without the ability to classify or quantify disease severity on the spike. Recently, image classification based on both presence and level of disease has been performed. Using a multi-year, multi-rater dataset captured using RGB cameras, researchers at AIMotion Bavaria were able to detect disease in the field and classify spikes into six severity levels by employing an EfficientNet-based neural network for image analysis. Images from multiple years improved the accuracy and decreased the root mean square error and in-field inter-rater reliability was lower than the model to raters (Rößle *et al*., 2023). Manual photography with commercial cameras requires additional human operators and cameras to increase the number of plots and images that can be collected, which limits phenotyping efforts for many large breeding and research programs.

By utilizing high-throughput phenotyping platforms, the scope of in-field disease detection can be expanded by reducing imaging time and labor. Images captured using a push cart were analyzed using a mask region convolutional neural network to detect FHB symptoms and assess severity on wheat spikes. The accuracies of wheat spike and disease detection were both high, and the pipeline was able to classify spikes into 15 severity stages with a 77% accuracy compared to ground truth (Su *et al*., 2020).

Previous studies have shown that detection of wheat spikes and FHB disease is possible in field conditions; however, low-throughput platforms for sensor deployment limits the scale of studies and the highly variable nature of FHB ground truth measurement makes model evaluation difficult (Rößle *et al*., 2023; Siou *et al*., 2014). Multiple raters are required to quantify the precision of human FHB measurements and create a reference standard for results. Thus, model efficacy should be determined by comparing accuracy against any individual human to determine if image analysis is a sufficient measurement replacement. Additional research is required to quantify multi-rater in-field and image based disease rating variation. This is necessary to empirically understand rater variation across phenotyping techniques, which would allow a more robust understanding of model outputs and human ratings, resulting in a deeper understanding and confidence in the model.

Advances in FHB image analysis and the development of a robust pipeline with demonstrated applications across years and camera configurations is needed to standardize phenotyping across research groups. The pipeline should be capable of precise disease quantification on single spikes to facilitate studies of host and pathogen interactions. In addition, the pipeline should be able to accurately aggregate plot disease scores for selections by breeders seeking to improve FHB resistance. This study describes a robust, neural network-based pipeline trained using images from multiple field locations in a single with the ability to quantify FHB in individual spikes across years. A comprehensive validation study was concurrently performed to assess pipeline results and quantify inter-rater reliability during field disease scoring and manual image annotations of FHB. Employing this disease assessment pipeline facilitates accurate and precise aggregate disease ratings for plots and opens novel avenues to elucidate host-pathogen interactions in the field through spike level resolution.

## Materials and Methods

### Field Design and FHB Inoculation

Wheat genotypes from the University of Minnesota wheat breeding program were grown in 2021 and 2022 at the University of Minnesota Agricultural Experiment Station in Saint Paul, MN (StP) and the Northwest Research and Outreach Center in Crookston, MN (Crk). Plants were grown in four row plots with each row being 1.5 m long, with 0.3 m row spacing, and 1.2 m alleys between plots. Plots in 2021 were planted in paired (two inner rows planted with the same genotype and two outer rows planted with winter wheat) and single (all four rows of the plot were planted with a different genotype) row configurations. In 2022, four replications of twenty genotypes were specifically planted in paired-row plots in StP and Crk for pipeline validation purposes. These genotypes represented a diverse range of spike colors, shapes, densities, and FHB disease responses.

In 2021, StP and Crk field trials were planted on May 12 and May 28, respectively. In 2022, StP was planted on May 10 and Crk on May 24. The field trials were inoculated with isolates of *F*. *graminearum* to promote FHB infection as previously described (Dill-Macky, 2003; Fuentes *et al*., 2005). Briefly, macroconidial inoculum of the pathogen was applied to the spikes when each plot was at anthesis using a CO_2_-powered backpack sprayer. A second inoculation was performed three to four days later to ensure that later tillers received inoculum at a time when they would be receptive to infection. In addition, paired row plots at StP and Crk were infected with grain spawn that was applied at the jointing stage and again seven days after jointing using previously described methods (Gilbert and Woods, 2006). To promote infection by *Fusarium*, the plots were irrigated with overhead sprinklers at or shortly before inoculation and continued through to the final disease assessment.

### Manual Field Ratings of FHB

Manual field rating of disease was performed in StP and Crk in 2022 by five raters, designated A-E. Disease severity was assessed as a plot aggregate. A score from 0-100% in 5% intervals was given by each rater based on the average FHB severity of spikes across a row. Prior to visual disease scoring, raters were shown a series of five diseased plots with a range of disease symptoms, including a susceptible check with >70% disease, to establish disease standards (Bock *et al*., 2022). The time required by each rater to manually score all the plots at a location was recorded. In StP, disease scoring was performed by all raters on July 18. In Crk, disease scoring by all raters was performed within a 24 hour time period between July 27 and 28. The rating dates were chosen to maximize disease progression, while minimizing the onset of natural plant senescence and were approximately 18-21 days after the final inoculations. Due to the confounding effect of natural plant senescence on disease assessment, plots designated as senesced were omitted from subsequent ratings and analysis. The same raters also completed manual disease annotations on images acquired during the 2022 season.

### Rover Based RGB Image Acquisition

Image acquisition was performed using an electric powered four-wheeled, semi-enclosed phenotyping rover designed, made, and supported by Mineral Earth Sciences LLC. (Figure S1a). The rover measured 2.1 m long and 3 m tall. The lateral wheel base was 1.3 m wide in StP and 1.2 m in Crk. A single person drove the rover with a remote control and also triggered image acquisitions. Imaging was controlled through a custom user interface through a Samsung Galaxy S20 smartphone that wirelessly connected to the rover through a MikroTik network router. Images from camera A and a Global Navigation Satellite System Real-Time Kinematic (GNSS-RTK) receiver allowed sub-10 cm precise registration of plots and captured images during operation. Images were stored on two internally mounted hard drives for data preservation. The rover accommodated eight Basler acA2500-20gc cameras for RGB image collection. In 2021, the rover was driven at 1.0 m/s. Images for model building from 2021 field trials were collected using side cameras C, D, G, H, I, and J (Figure S1b). After preliminary testing in 2021 to determine the optimal camera height, angle, and resolution for single spike imaging, cameras C and D were reconfigured for 2022 (Figure S1c). These cameras were mounted at 125 cm within the rover enclosure and oriented at 39 degrees to image plants between 30 cm and 112 cm in height. In addition, the rover driving speed was reduced to 0.5 m/s in 2022 to further improve image quality. The camera configuration change and reduced rover speed in 2022 facilitated high resolution imaging of spikes.

Imaging was performed once a week from emergence to heading, and twice a week from the onset of disease symptoms through senescence. In 2021, image collection was performed on single and pair-row plots from June 11 to August 10 in StP and June 16 to August 3 in Crk. In 2022, images from paired-row plots were captured on ten different dates in StP from June 17 to July 25 and six different days in Crk from June 23 to August 3. The rover captured images automatically at eight frames/second. In 2022, it took an average of 588 seconds to image 80 plots at each location. An average of 37643 images were taken during each imaging session from all eight cameras. A single plot took approximately 7 seconds to complete and resulted in 471 images from all cameras, which is approximately 59 images per camera per plot. After images were registered to plots and the edges of plots excluded to remove any edge effects, there were on average 22 images captured per camera per plot.

### RGB image processing

Images from each plot were selected, aligned, and overlaid using GNSS-RTK coordinates. Plot boundaries were drawn and future images were registered to these predetermined plots based on location in subsequent scans. In addition, images registered outside plot boundaries, including alleys and edges of the field, were omitted from downstream analysis. In 2021, images from cameras C, D, G, H, I, and J were randomly sampled between June 22 and August 3 for model building (Figure S1d). In 2022, a subset of eight images, four each from cameras C and D, were sampled across the plot to minimize repeated images of spikes in pipeline assessment (Figure S1e, Figure S2).

### Modeling workflow

An automatic image analysis pipeline was developed to output FHB disease levels per spike using images collected by the phenotyping rover (Figure 1). Four models were developed that detected and segmented wheat spikes in the field, segmented FHB disease on the spikes, and determined spike gradability (image quality). For each model, a random sample of images from paired and/or single row plots were used from 2021 collected images. Images acquired from the 2022 field trials were not used in any model training, but instead were utilized for overall pipeline assessment and cross-year model validation. All image annotations were done by the raters using an internal Google data labeling platform.

**Figure 1.**
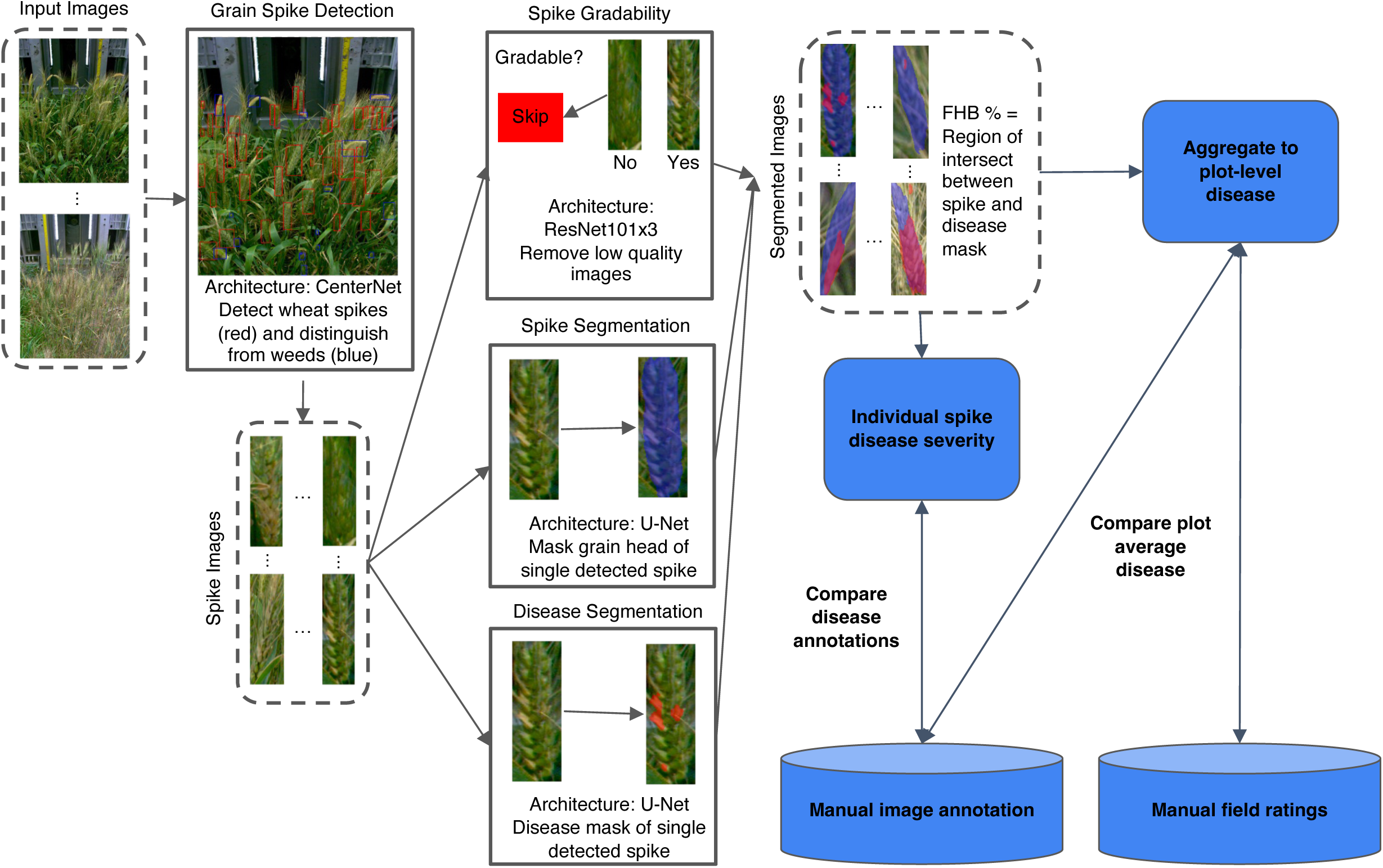
Overview of the FHB image analysis pipeline, associated model architecture, and comparisons used for model validation. The pipeline was made up of four models. The grain spike detection model identified wheat spikes in complex field images. The spike and disease segmentation models segment entire unobscured wheat spikes and the portions exhibiting FHB symptoms, respectively. Disease severity was quantified as the region of intersection between the segmented spike and disease. The spike gradability model was used to assess image quality before plot level disease aggregation. The pipeline was validated by 1) comparing the percentage of disease on individual spike images with manual image annotations done by raters, 2) comparing a plot aggregate disease score based on gradable spikes annotated by the pipeline with plot aggregate of large scale manual annotation of spike images by humans, and 3) comparing a plot aggregate disease score based on gradable spikes annotated by the pipeline with manual field rating plot disease scores.

#### Spike Detection

A spike detection model was developed to identify wheat spikes in complex images from variable field conditions (Figure 1). Labeled examples from 2021 field trials spanning multiple dates in StP and Crk were used to capture spikes at different growth stages. Images were randomly selected from plots throughout the field to capture diverse spike morphologies. Images from side cameras I, J, C, D, G, and H in 2021 were used for model development (Figure S1b). A collection of 2048 images were randomly split into 90% training and 10% evaluation sets. Spikes were categorized into two classes: “wheat spike” or “other”. The “other” class consisted of weeds with similar characteristics as wheat spikes, such as foxtail (*Setaria* species). This explicitly trained the model to learn the difference between wheat and non-wheat spikes to reduce false positives. Input images were resized to a resolution of 1024 x 1024 pixels and roughly 80% of images that contained no labeled wheat spikes were removed to ensure that images from before heading did not overwhelm the model during training. The CenterNet architecture with an Hourglass-104 backbone was used for object detection and initialized with weights from models trained to detect wheat spikes from images collected in Mexico. This model identifies objects as a triplet using information from the top-left, bottom-right, and central regions to improve precision and recall (Duan *et al*., 2019).

#### Spike Segmentation

To determine total spike area for subsequent disease annotation, a spike segmentation model was trained (Figure 1). The labels for the model were obtained by randomly sampling 1000 images from dates after heading, running the spike detection model on each image to identify wheat spikes, and then randomly sampling at most five spikes from each image for labeling by raters. This ensured a wide variety of spike morphologies were used in developing the model. Each training example consisted of a single detected spike, with a mask drawn over the wheat spike. In the event that another spike was present within the bounding box due to close proximity, the center most spike was labeled with a mask. Images were resized to 128 x 128 pixels prior to training and any images lacking masks (i.e. potentially representing false positives from the detection model) were dropped. The U-Net deep learning architecture was used for semantic segmentation (Ronneberger *et al*., 2015). The dataset for the model consisted of 4974 spikes which were randomly split into 90% training and 10% evaluation sets.

#### Wheat FHB Segmentation

To develop the FHB segmentation model, the five raters were provided with randomly sampled full images containing spikes within a plot captured from camera C or G in 2021. A bounding box was placed around a single detected spike for disease annotation in order to provide plot level context to the rater. At most, 5 spikes per image were labeled (Figure 1). A total of 1208 images were annotated by the five raters, 313 from July 22, 332 from July 27, and 563 from August 3. The dates that the images were sampled from were the dates closest to when in-field ratings were performed to ensure that sufficient levels of FHB symptoms were present. The number of images for annotation of FHB symptoms on the spikes was evenly spread across the five raters. The raters drew a polygon around the symptomatic region(s) on the spike to label the disease for model training. Cropped spikes with manually annotated disease masks were used to train the U-Net architecture for disease segmentation (Ronneberger *et al*., 2015). Training images were resized to 256 x 256 pixels and were randomly split into 80% training and 20% evaluation sets.

#### Spike Gradability

A wheat specific gradability model was trained to ensure high image quality of spikes used for final disease assessment (Figure 1). A total of 3072 annotated spike examples were used for model training. Images were taken from the same dates used for disease annotation. There were 854 images used from July 22, 2021, 788 from July 27, 2021, and 1430 from August 3, 2021. The raters “skipped” spike images of low quality when performing their FHB disease segmentation labeling. The spikes were assigned an “ungradable” label (label ID = 1) if the spike was skipped, otherwise the image was deemed “gradable” (label ID = 0). Of the 3072 sample spikes, 1376 (44.8%) were found to be gradable and 1696 (55.2%) were ungradable. Although a high percentage of spikes were deemed ungradable, this was expected as the image is capturing many heads in the plot, but not all of them are in the optimal focal range of the camera. The frame rate of the camera and number of times each spike was represented in subsequent images as the rover moved through the field gives us confidence that the majority of non-occluded spikes will be deemed gradable in at least one image. Gradable spikes were typically in focus with good color contrast for raters to identify and label FHB symptoms. Ungradable spikes were often blurry, small, over or under exposed, occluded, a mistaken weed species, or marked with spray paint for field marking purposes. Gradability sample images were fed into a deep learning model based on ResNet101×3 to maximize accuracy gained from increased layer depth (He *et al*., 2015). Data was randomly split into 80% training and 20% evaluation sets. The training examples had 1109 gradable and 1333 ungradable images and the evaluation set had 267 gradable and 363 ungradable spike examples.

#### FHB Severity Calculation

The calculation of FHB severity per spike was conducted by determining the ratio of disease-afflicted pixels to the total number of pixels constituting the wheat spike (Figure 1). Only disease pixels that intersected with the spike segmentation pixels were considered in this computation. Any instances of disease presence occurring outside the spike, potentially attributable to background interference, like the presence of another head or unrelated discoloration that might resemble disease, were excluded from this calculation by using spike segmentation boundaries.

### Manual Image Annotation

To assess the precision and accuracy of the FHB automatic image analysis pipeline, manual image annotation was performed on images captured during the 2022 field season and compared against model inference results. Images were selected from Crk on July 28 and StP on July 18 to align with manual field rating dates. Four of the five raters used to annotate images for model building in 2021 were retained for model evaluation efforts in 2022. These raters will be referred to as raters A-D. A new fifth rater, rater E, was added in 2022 for additional validation efforts. Rater expertise ranged from complete novice to experts with more than a decade of in-field FHB disease scoring experience.

Two subsets of images for manual disease annotation were created. The first subset had 8184 distinct spike images for large scale annotation in order to judge rater skip rates and obtain enough disease scores to calculate plot averages. To determine the number of spikes to annotate, a dynamic sampling strategy was employed based on 2021 model inferences. This sampling strategy was used to minimize required labeling time. Taking into account that FHB infection is typically not uniform across a plot, the standard deviation of disease severity for spikes within each plot was calculated. It is expected that plots with around 50% disease severity will exhibit the highest standard deviation compared to lower and higher disease levels. Thus, plots with low and high disease levels would require fewer labels to capture the variation within a plot compared to plots with a moderate level of disease. With a specified FHB error tolerance of 2c = 10%, where c = 5%, the required number of labels was determined as (standard deviation / c)^2^. The total number of labels sent for labeling was initially calculated assuming 70% gradable spikes and a 99% probability of reaching the desired number of required labels per plot. This number was rounded up to distribute an equal number of labels to each rater. To account for potential incorrect labels, the required number of labels had a factor of safety of 1.5. Each rater was given a unique random sample of spikes from the 8184 spikes distributed across StP and Crk paired-row plots. Raters A-E first assessed image quality and then determined if images were gradable or ungradable. Images deemed ungradable were skipped. Each image was then manually annotated for areas showing FHB symptoms using a Google internal data labeling platform with a freeform shape tool.

From the gradable spikes in the large scale annotation image subset, 200 spikes were selected to assemble an inter-rater reliability annotation image dataset. Each of these images was annotated by all five raters. To assemble the dataset, labeled large scale annotation spikes were binned into discrete categories based on disease severity, from 0-10% to 80+% in 10% increments. Spikes were randomly selected from each bin to ensure all disease scores were represented in the inter-rater reliability analysis. Thirty spikes were sampled from the [0-10] and [10-20] classes because low disease percentages occurred more frequently in the dataset. Twenty spikes were selected from each of the remaining categories. Six spikes were selected from each rater for the 30-label categories, and four spikes were selected from each rater for the 20-label categories. Images from StP and Crk were sampled evenly.

### Statistical analysis

Data analysis and visualization of manual field ratings, manual image annotation, and pipeline image annotation results was performed in R version 4.3.1 within R Studio version 1.4.1717 using the ggplot2 and tidyverse packages (Wickham, 2016; Wickham *et al*., 2019; RStudio Team, 2020; R Core Team, 2022). Linear regression and related functions were performed using the R lme4 package (Bates *et al*., 2015). Intraclass correlations were performed using the R irr package (Gamer *et al*., 2019).

## Results

### Manual field rating agreement between raters is affected by disease severity and experience

FHB infection can be quite variable among wheat plants within a row, making visual assessments of plot aggregate disease severity difficult. To establish a minimum correlation threshold for evaluating our FHB image analysis pipeline, five raters scored aggregate FHB severities for plots in StP and Crk in 2022. In both locations, 80 plots were assessed for FHB levels. The phenotyping rover was run at these locations on these dates to pair subsequent images used for manual (done by raters) and pipeline image annotation (model output) with field ground truth. In Crk and StP, the average disease severity of visual manual in-field ratings was 21.4% and 37.7%, respectively (Figure 2a). Of the five raters, raters A-D had very similar disease scores and standard deviations, which shows the variation of FHB severity between plots could be accurately discerned. The average disease scores for raters A-D were 18.4 to 26.1% (standard deviation of 12.8 to 16.3%) in Crk and 33.1 to 36.3% in StP (standard deviation of 13.9 to 21.7%). In Crk, rater E had the lowest average disease score of 12.9%, while in StP rater E had the highest average disease score of 48.9%. In both Crk and StP, rater E had the lowest standard deviations of 9.4% and 10.6%, respectively.

**Figure 2.**
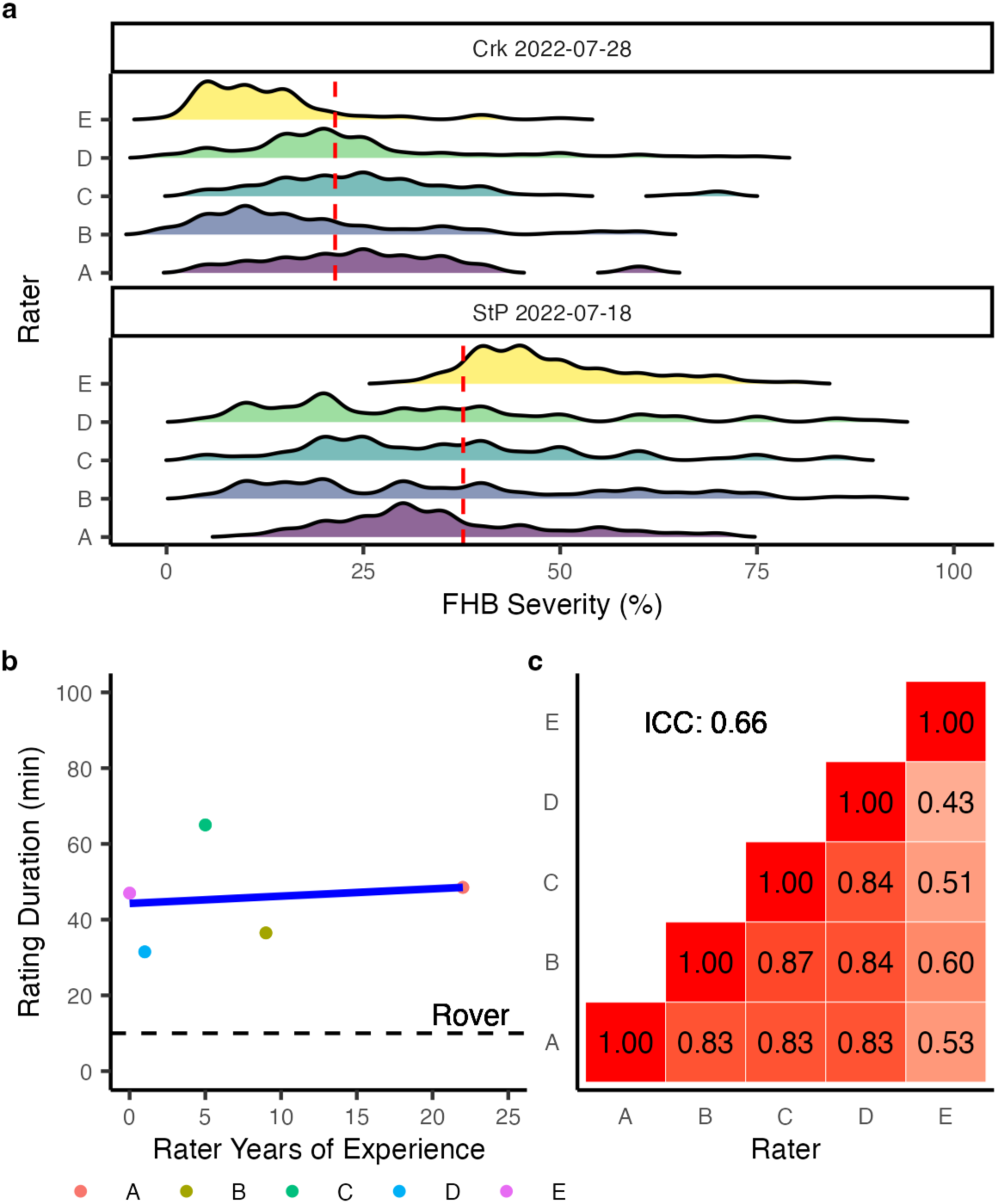
FHB severity, disease scoring duration, and rater agreement for manual field ratings conducted at Crookston (Crk) and Saint Paul (StP) in 2022. a. Distribution of plot level disease severity scores from each rater for Crk on July 28 and StP on July 18. The mean disease severity for each location is shown by the red dashed line. b. The relationship between rater experience and rating duration. The regression line between experience and time is shown in blue. The dashed black line is the average imaging time for the phenotyping rover for all the plots. c. Pairwise Pearson correlation coefficients and intraclass correlation (ICC) values between rater in-field disease scores.

The research experience of the raters working on FHB ranged from 0 to 22 years (Figure 2b). Rater E had the least experience, with 2022 being their first summer of disease rating. Rater A had the most experience with 22 years of FHB scoring. The average duration that each rater took to assess FHB on 80 plots at the two locations ranged from 23 to 80 minutes. On average across all the raters, it took 44 minutes to manually rate the plots in Crk and StP. For the limited sample size of raters in this study, there was not a statistically significant relationship between FHB experience and rating duration (*p = 0.77, R² = 0.01*). In contrast, the phenotyping rover took about ten minutes, approximately 20% of the time required for a single person to visually rate the same plots in the field.

The agreement of the five raters on in-field plot level disease scoring was influenced by location-specific disease severity and rater experience. When combining the in-field disease ratings from both locations, pairwise Pearson correlations between any two individual raters ranged from 0.43 to 0.87 (Figure 2c, Table 1). The least experienced rater, rater E, had on average a 0.51 correlation with the other raters which was the lowest correlation. In contrast, raters A-D had at least one summer of FHB rating experience and correlated highly with one another with an average 0.84 correlation. An intraclass correlation was calculated to understand how raters’ in-field assessments correlated when considering them as a group instead of in pairwise fashion. The intraclass correlation among all raters across both locations combined was 0.66, with a 95% confidence interval of 0.60 to 0.73 (Table 1). When looking at the correlation of raters in each rating location, StP had an average Pearson correlation of 0.69 and an intraclass correlation of 0.51, with a 95% confidence interval of 0.40 to 0.62. The Crk location had an average Pearson correlation of 0.87 and an intraclass correlation of 0.70, with a 95% confidence interval of 0.62 to 0.78. This indicates location influenced the correlations of FHB disease ratings of the raters, with the Crk location having closer in-field disease scores between the five raters.

**Table 1.** Plot level FHB correlations and intraclass correlation (ICC) between the average disease level from the five in-field visual raters, the plot average disease level from imaged spikes manually annotated in the large scale annotation image dataset, and the plot average disease level based on all the spike images automatically annotated by the developed FHB pipeline.

|  | Manual Field Rating | Large Scale Manual Image Annotation |
| --- | --- | --- |
| Manual Field Rating | ICC = 0.66<br>r = 0.43-0.87 | r = 0.74 |
| Automatic Image Annotation | r = 0.85 | r = 0.75 |

### Relationship between in-field plot averages, manual image annotations plot averages, and model output of individual spike disease scores

To compare the precision and accuracy of manual image annotations to in-field ratings for assessing disease severity, two subsets of images were taken, one from Crk images taken on July 28, 2022 and one from StP images taken on July 18, 2022 (Figure 3). For both datasets, it took raters approximately 30 seconds to annotate disease on each spike. The large scale manual annotation image dataset consisted of 8184 images and was used to evaluate gradability and FHB disease severity per spike. Within this dataset, 54% of the spike images were skipped by the annotators, which classified the spike as ungradable. Between locations, an average of 46% of the spike images from Crk and 59% of the spike images from StP were labeled as ungradable by the five raters. The skip rates of spikes, or ungradable classification, among raters ranged from 32% by rater B to 84% by rater A. This indicates a strong subjectivity among raters for what constitutes an unambiguous or high enough resolution image of a spike that can be reliably rated for FHB symptoms. Of the total 8184 spike images, 4526 were annotated for disease by the raters. Of the 4526 spike images that were annotated, raters B, D, and E annotated an average of 1177 images, while raters A and C annotated 497 images on average. The average FHB percent annotated on spikes by each rater was 13.11%; rater D annotated the lowest average FHB percent per spike at 11.7%, while rater C annotated the highest at 15.2% (Figure S3). This indicates that raters annotating high resolution images of wheat spikes for FHB severity would have a high level of agreement.

**Figure 3.**
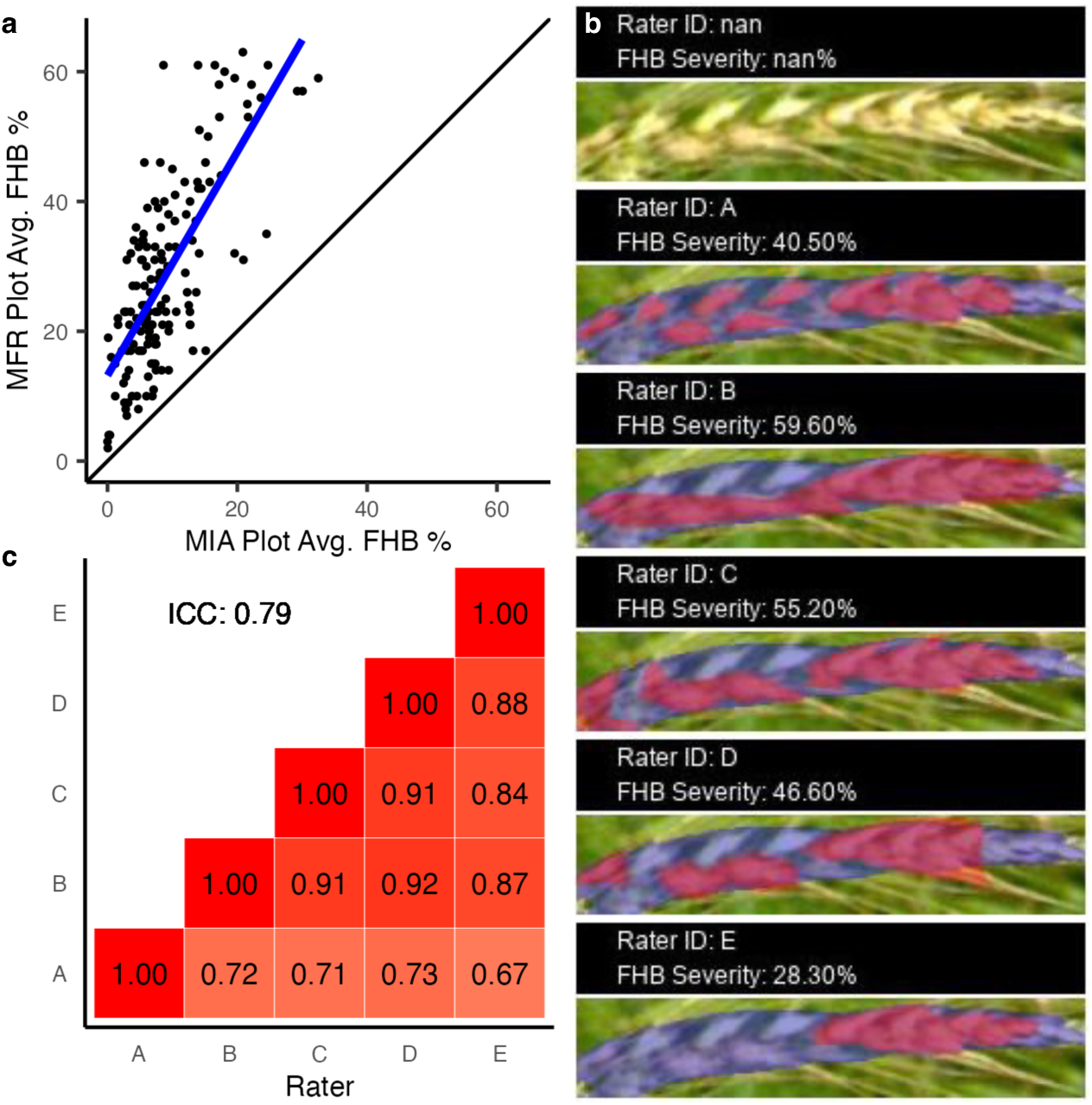
Manual image annotations (MIA) of FHB severity correlations between raters and in-field FHB ratings. a. Relationship between the manual in-field rating (MFR) of FHB plot average of all five raters and the plot average of FHB percent of spikes from the large scale manual image annotation (MIA) dataset. The black line indicates a one to one relationship. The regression line between the manual image annotation and manual in-field rating averages per plot is shown in blue. The Pearson correlation coefficient of this comparison is 0.74. b. Repeated images of the same representative image from the 200 inter-rater reliability annotation dataset. The image was selected to represent the average coefficient of variation of the inter-rater reliability image annotation set. The mask of the spike segmentation is shown in purple and the FHB disease annotations on the spike by each of the five raters is shown in red. The level of FHB severity annotated by each rater is also shown. c. Pairwise Pearson correlation coefficients and intraclass correlation (ICC) between raters for the 200 inter-rater reliability annotation dataset.

Using the large scale manual image annotation dataset, the percentage of disease of all gradable spikes (from all five annotators) from the same plot were averaged together to get an aggregate disease score (Figure S4). This was compared with disease scores from in-field ratings averaged from all five raters to determine rating differences between spike images and in-field ratings. The average plot disease severity from manual image annotations across the five raters was 9.7%, while the average in-field plot disease severity was 28.7%. This shows either an over-inflation by raters in the field or a deflation of disease scoring when raters are looking at individual spike images. There was a 0.74 Pearson correlation between the large scale manual image annotation and field plot averages (Table 1). Pairwise correlations between plot averages rated in the field by a rater and the subsequent plot averages calculated from the spikes they annotated in the large scale manual image annotation ranged from 0.75 to 0.81 for raters A-D. Rater E, the least experienced rater, had a 0.34 correlation between their in-field ratings and plot averages calculated from the spikes they annotated in the large scale manual image annotation dataset.

The 200 images used for inter-rater reliability assessments were sampled across the disease spectrum from spikes determined to be gradable during the large scale manual image annotation evaluations. This meant that a rater determined the spike gradable; therefore the image of the spike would be considered of sufficient quality to have all raters annotate it for FHB to determine how raters compare in their assessments when annotating the same exact spike image. Each spike image from this 200 image dataset was annotated for disease by all five raters (Figure 3b). The relative distribution of disease across the disease spectrum was similar among all raters (Figure S5). Notably, there were less spikes graded as moderately diseased, approximately 50 to 70% disease, by the five raters than was expected in the theoretical disease distribution. Likewise, there were also more spikes annotated between 20-30% by raters than expected in the theoretical distribution.

Annotations from the inter-rater reliability image dataset showed that, while the exact disease annotations varied between raters, the general location and distribution of disease annotations on spike images was consistent among all five raters (Figure 3b). A coefficient of variation was calculated to quantify the spread of disease percentages for each spike annotated by all of the raters. The average coefficient of variation for the 200 spikes was 0.38, which indicated a moderately low dispersion around the mean FHB percentage. The Pearson correlations and intraclass correlation were calculated to test the relatedness among raters for the inter-rater reliability annotation images (Figure 3c). Pairwise agreement was generally high, ranging from 0.67 to 0.92 (Table 2). The most experienced rater, rater A, had the lowest correlations with the other raters. Rater E, the least experienced rater in this study, had comparable FHB annotation correlations with the more experienced D, C, and B raters. Overall agreement among the raters, as measured by intraclass correlation, was 0.79, which is 0.13 higher than the intraclass correlation of manual in-field ratings (Table 1). Based on these results, raters correlate better when looking at high quality images of spikes than when rating whole plots in the field.

**Table 2.** Single spike image FHB correlations and intraclass correlation (ICC) between the 200 image inter-rater reliability image dataset, the large scale manual image annotation dataset, and the automatic image annotation of FHB on spike images by the FHB image analysis pipeline.

|  | Inter-Rater Reliability<br>Annotations (200 images) | Automatic Image<br>Annotation (model) |
| --- | --- | --- |
| Inter-Rater Reliability<br>Annotations (200 images) | ICC = 0.79<br>r = 0.67-0.92 | ICC = 0.75<br>r = 0.63-0.83 |
| Large Scale Manual Image<br>Annotation | N/A | r = 0.70 |

### The automatic image analysis pipeline output correlates with in-field and manual plot averages

The automatic image analysis pipeline was trained using 2021 data and run on images taken from Crk on July 28, 2022 and StP on July 18, 2022 for comparison with the manual in-field ratings and the large scale manual image annotation results. It took 4.75 hours to run the complete pipeline on all images collected by the rover on these dates. Disease inferences outputted by the pipeline were examined at a plot and individual spike level. The automatic image analysis pipeline was run on images and a total of 109501 spikes were detected using the head detection model (Figure S6a), 61658 spikes from StP and 47843 spikes from Crk. For each spike, the percentage of FHB and a gradability probability were determined by the spike segmentation, disease segmentation, and spike gradability models. Prior to filtering spikes by gradability probability, the average spike FHB percentage in StP was 28.8% and the average in Crk was 6.1%. The distribution of spike FHB percentages skewed towards the lower end of the disease spectrum (Figure S6b). The gradability threshold was based on a classification scale from 1.00 (ungradable) to 0.00 (gradable). Based on this classification model, images with a gradability probability <= 0.50 were classified as gradable and kept for further analysis. The number of retained spikes and the correlation with in-field and the large scale manual image annotations stabilized at a gradability probability <= 0.50 (Figure 4a). After applying the gradability probability filter, across the 80 plots at each location 8080 spike images were retained, 4239 from StP and 3841 from Crk (Figure S6c). Again, the distribution of spike FHB percentages skewed towards the lower end of the disease spectrum (Figure S6d). The average percent FHB in StP after filtering was 21.7%, which is lower than before filtering ungradable spikes, and the average Crk disease was 6.8%, which is similar to before the gradability filter was applied.

**Figure 4.**
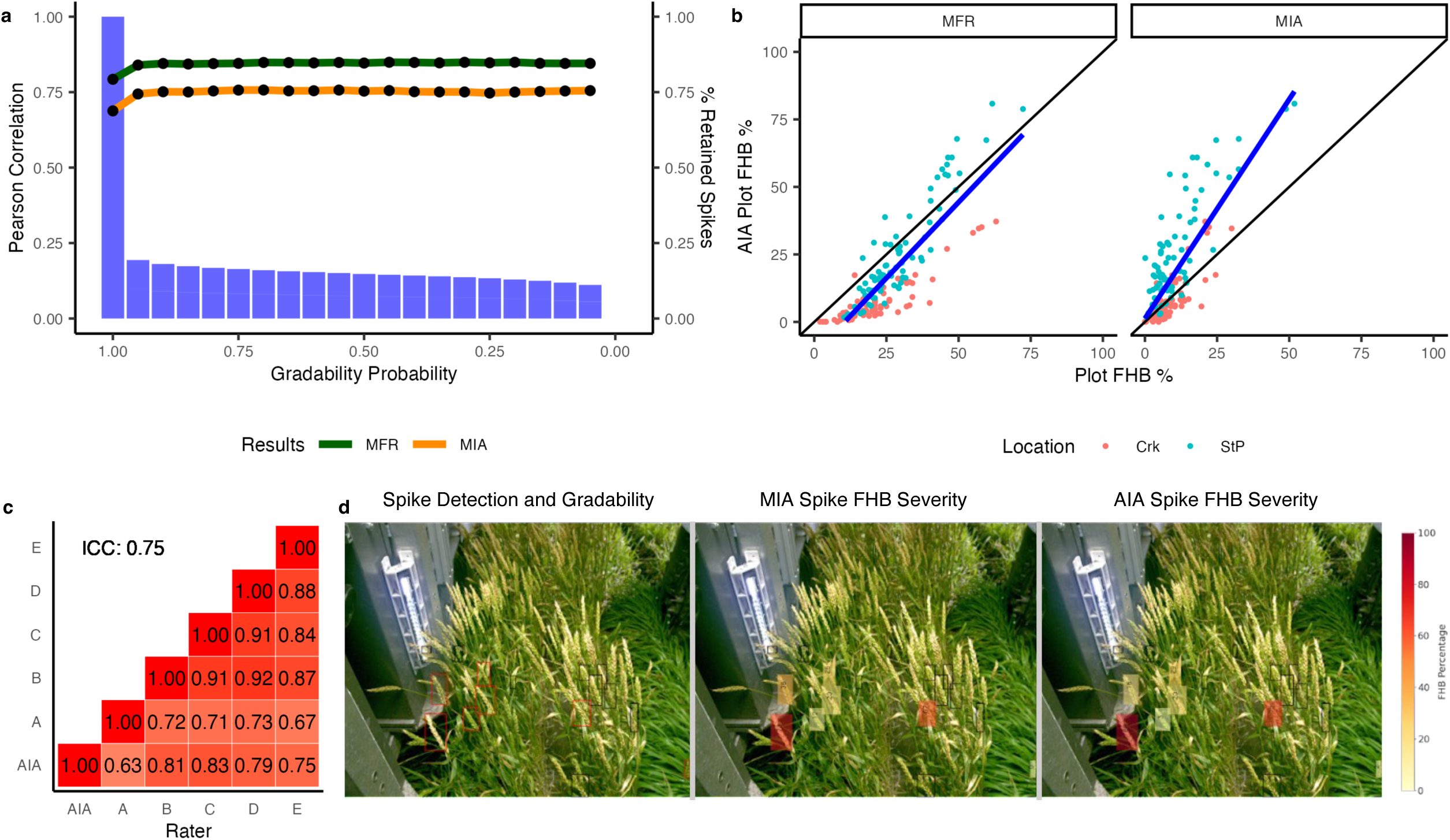
The output and validation of the automatic image analysis pipeline results. a. The effect of the gradability model threshold on the number of retained spikes and the correlation to manual in-field ratings (MFR) and the large scale manual image annotation (MIA) plot averages. b. The automatic image analysis (AIA), which is model output from all spikes in all images, plot averages plotted against the manual in-field ratings (MFR) and the large scale manual image annotation (MIA) plot averages. The regression lines for each plot are shown in blue. c. Pairwise Pearson correlation coefficients and intraclass correlation (ICC) between raters and the automatic image analysis pipeline for the 200 inter-rater reliability image dataset. d. A representative example of model spike detection, large scale manual image annotation FHB percentage, and automatic image analysis pipeline inference results. The gradable spikes are shown with red bounding boxes and the black bounding boxes marked the ungradable spikes. The color of the filled boxes shows the FHB percentage for the gradable spikes calculated by each method.

After filtering for gradability, there was a mean of 52 spikes and a median 47 spikes per plot. Plot level disease averages were calculated from the output of the automatic image analysis pipeline results and compared to manual in-field rating plot averages that were calculated by averaging all five rater disease scores (Figure 4b). A Pearson correlation of 0.85 was observed between the model results from the automatic image analysis pipeline and the manual in-field rating results (Figure 4a/b, Table 1). The Pearson correlations within separate locations were slightly higher in StP than in Crk, 0.92 and 0.88, respectively. The plot averages from the large scale manual image annotation image dataset, calculated by averaging spike FHB percentages from all five raters within a plot, were also compared to the automatic image analysis pipeline model inferences (Figure 4b). The Pearson correlation between the automatic image analysis pipeline and the large scale manual image annotations was 0.75 (Figure 4a/b, Table 1), and 0.82 and 0.83 in StP and Crk, respectively.

To determine if the image analysis pipeline could accurately determine FHB percentage of individual spikes, the automatic image analysis pipeline disease inferences were compared to the inter-rater reliability dataset (200 spike images). As the spikes in this dataset had already been classified as gradable by the raters, no gradability filtering was applied. When the automatic image annotation pipeline was used to inference FHB severity on spikes from the inter-rater reliability annotation subset, the intraclass correlation between model results and all rater annotations was 0.75 (Figure 4c, Table 2). The Pearson correlation with individual rater annotations ranged from 0.63 to 0.83 (Table 2). The automatic image analysis on the inter-rater reliability dataset correlated with all raters equally or slightly less than pairwise correlations among raters B-E, and exceeded rater A correlations with the other raters using the 200 image dataset (Figure 4c). Interestingly the performance of the automatic image analysis pipeline was influenced by location. In StP, the intraclass correlation between the pipeline output and raters using the inter-rater reliability dataset was 0.67, with the Pearson correlation for individual raters ranging from 0.58 to 0.82. Inferences on spikes from Crk had an intraclass correlation of 0.84, with the Pearson correlation ranging from 0.75 to 0.94. In all comparisons the automatic image analysis pipeline FHB severity inferences had the lowest agreement with rater A.

The performance of the automatic image annotation pipeline on spike images from the large scale manual annotation image dataset was compared to manual annotations on the large scale manual image dataset (Figure 4d). Unlike the inter-rater reliability image dataset, raters did not annotate the same spikes so no intraclass correlations could be calculated. There was a 0.70 Pearson correlation between the pipeline and rater disease annotation results, with individual model-rater pairs having correlations between 0.65 and 0.82 (Table 2). Images from StP had a lower overall correlation of 0.70 between model and raters compared to Crk, which had a higher correlation of 0.89. The average disease per spike for StP was 36.1% from the automatic image analysis and 13.6% from the raters and in Crk it was 10.9% and 11.6%, respectively (Figure S7).

## Discussion

### Image annotation is faster, more precise, and more accurate than manual field ratings

Manual disease rating is tedious, slow, and results are often subjective and biased by human error (Araus *et al*., 2018). Previous research has shown that visual ratings may be influenced by overestimation of disease at low disease severities, preferred rating values, and confounding host characteristics (Bock *et al*., 2020). When assessing the accuracy and precision of our FHB ratings, Pearson correlations and intraclass correlation values were used to assess relative rater agreement, rather than using mean absolute error due to the lack of a gold-standard FHB score available for reference. Instead, we prioritized accurately capturing variance in disease levels, similar to how breeders will often use reference varieties to compare FHB reaction, rather than using absolute disease severity numbers. We found that, while manual in-field rating correlations between experienced raters were high for FHB, the inexperienced rater with no prior FHB scoring experience had extremely low correlations with other raters. Seasonal, field research teams are often made up of new, inexperienced raters. Employee retention and training represents a significant barrier to researchers studying FHB. Combined with the aforementioned limitations-cost, speed, tedious nature, and subjectivity of rating FHB disease ratings can quickly be compromised and error prone which will reduce the heritability of disease resistance.

Image annotations by raters and the developed automatic imaging analysis pipeline were more precise than field disease scores. The agreement of raters across the manual image annotation datasets was higher than rater agreement of FHB visual disease scoring in the field, including between both new and experienced raters. Moderately strong correlations between plot averages from spikes in the manual image annotation dataset and in-field ratings demonstrated that image analysis is an appropriate and effective method for in-field FHB quantification. Although more precise and accurate than in-field ratings, annotating individual spikes from images is too time and labor intensive to be a viable replacement for disease assessment. As a limited number of spikes could be reasonably be manually annotated in each season, plot averages would be determined by a small sample of spikes which might not capture the true FHB severity of the plot. By taking the manual image annotation results we developed an automated image annotation pipeline that was able to have high accuracy and precision in FHB severity rating and also be able to process images with little labor and maximize the number of spikes evaluated. The automatic image annotation inference results were generated about two to three times faster than manual annotations and parallelization would greatly improve the run time of the pipeline on large image datasets.

High-throughput phenotyping methods also increase the speed of data collection across traits and species (Tanger *et al*., 2017; Sun *et al*., 2017). The in-field rating method used in this study was chosen to maximize speed for comparison against the rover, yet visual disease rating was about five times slower than acquiring images with the rover no matter the experience level of the rater. More objective disease quantification methods that might improve inter-rater reliability would require even greater time and labor investments (Stack and McMullen, 1998). These time constraints limit the number of genotypes and environments that can be evaluated for FHB in any given year.

### Automatic image analysis pipeline can accurately quantify FHB severity on a spike and per plot basis

Previous FHB deep learning models have had success in spike and disease detection and classification (Hong *et al*., 2022; Su *et al*., 2020; Rößle *et al*., 2023). To push the research forward requires efficient disease quantification on individual spikes at scale. Using the high-throughput rover and our automatic image analysis pipeline for FHB severity, we demonstrated the ability to accurately and efficiently quantify FHB severity on an individual spike and plot levels. The automatic image analysis pipeline inferences for single spike images displayed moderate to high accuracy when compared to manual image annotation results. When comparing the same spikes annotated by the automatic image analysis pipeline and by raters in the inter-rater reliability image dataset, the intraclass correlation showed little change when adding pipeline FHB severity inferences with rater annotation results together (Table 2). The range of pairwise correlations between the pipeline FHB severity ratings and individual raters was also within the range of correlations among rater pairs for all manual image annotation datasets.

Some differences between FHB severity pipeline and rater disease scores may be due to individual human tendencies during image annotations. For example, there was a slight deviation between theoretical FHB severity and human annotations for inter-rater reliability images (Figure S5). In particular, fewer spikes were rated as moderately diseased (∼50-70%) than expected based on pipeline inferences, but more spikes were annotated at the ends of the FHB disease spectrum (<25% and >75%). This may be due to a tendency of raters to rate small individual areas of disease or mark an entire spike as diseased, but be less thorough when multiple, variable areas of disease symptoms were present. It was also observed in annotations of the manual annotation image sets that raters had their own tendencies during disease annotations. For example, rater A had the lowest correlations in inter-rater reliability images and also had highest skip rates during large scale manual image annotations. There were multiple spikes marked as 0% disease by rater A that the model and other raters annotated as greater than 75%. Due to the difficulty and scope of phenotyping FHB, a definitive gold standard disease score is difficult to obtain in these cases. These results demonstrate that even when controlling for confounding factors in manual image analysis, it is difficult to completely remove human bias in disease phenotyping, no matter the nature of the task.

The imaging location also had an impact on the automatic image analysis pipeline results. The intraclass correlation between raters and the pipeline for the 200 image inter-rater reliability dataset was 0.84 in Crk, compared to 0.67 in StP. Similarly, the correlation between the pipeline and the large scale manual image annotation spike dataset was 0.70 in StP and 0.89 in Crk. Furthermore, the lowest pairwise correlations between manual rater and the pipeline inferences were always found in StP. Plant-pathogen interactions may partly explain differences between locations. Severity of FHB infection and rate of wheat flowering and maturation is heavily influenced by environmental factors (Moreno-Amores *et al*., 2020). The U-Net model trained to segment disease has shown good performance in complex backgrounds, but location impact on morphology and phenology may still affect model performance (Liu and Wang, 2021). In addition, spike images from StP have a more muted green saturation than Crk images, which could seem overexposed in some cases (Figure S8). This may have caused the FHB detection model to overestimate at StP and increase the calculated single spike disease percentage.

One strength of high-throughput phenotyping is the ability to score many spikes within a field to obtain accurate plot disease estimates. The FHB severity pipeline was able to predict plot aggregate disease ratings with high accuracy. Correlations between the pipeline and manual image annotation plot averages were equivalent to correlations between in-field and manual image annotation plot averages. Unlike on an individual spike basis, location was not seen to impact relatedness between the pipeline and in-field or manual image annotation plot averages. This shows that over a large random sample of images, the pipeline is able to accurately quantify aggregate FHB severities regardless of differences in locations. The automatic image analysis pipeline plot averages had a higher correlation with in-field ratings than manual image annotation scores. We hypothesize this was due to the tendency of the model to inflate disease on certain images from StP, conflated with the tendency of human raters to overestimate disease during field scoring. The average FHB severity was rated higher in the in-field ratings than the manual image analysis. This coincides with previous research that showed disease levels on individual spikes tended to skew lower during image analysis (Su *et al*., 2020). Therefore, higher image disease scores would shift the pipeline plot averages to be more similar to manual in-field rating results. Conversely, this could be viewed as rater image annotations being too conservative compared to in-field and the pipeline inference results.

Another reason for the discrepancy between automatic and manual image annotation results may be due to the gradability classification procedure. For plot aggregate disease scores, a gradability threshold <= 0.50 was used to filter ungradable spike images. For comparison to manually annotated images, gradability was not used to preserve manual image annotations performed by raters, and allow a one-to-one comparison between the pipeline and raters to be made. However, a drawback of this may be that even though a single rater determined a spike image gradable, not all the others raters might agree. Also, as classifications done by raters were used to train the gradability models, the pipeline might still not be optimized to annotate images with a gradability >0.50. This could alter correlations between the automatic and manual image annotation analysis. As the model is refined using 2022 images, we expect the number of gradable spikes to increase, which may subsequently improve correlations to rater manual image annotation results.

### The pipeline is robust across years, camera angles, and locations, however additional refinement may be required to optimize across environments

Along with being more reliable than in-field disease ratings and more efficient than manual image annotations, these results demonstrate that the developed FHB severity automatic image analysis pipeline is broadly relevant across a range of real-world research applications. One limitation of current FHB research is the lack of a consistent method for disease assessment. Different lab groups use different methods to score FHB, which makes comparing and consolidating results difficult. The automatic image analysis pipeline demonstrated success calculating FHB severity on spikes and plots across multiple years and locations. Furthermore, the 2021 images used to train the different models used in the pipeline were captured with a different camera arrangement than what was used to collect images in 2022 (Figure S1d/e). The ability of our models to extrapolate disease beyond its training configuration increases the potential for this tool to be useful for researchers without access to the phenotyping rover. However, we found that 2022 spike images were smaller than 2021 training images, which may have impacted the gradability threshold. Therefore, to maximize the number of usable spikes from the pipeline, some fine-tuning may be required for new camera orientations and imaging conditions.

To further test the ability of our model to provide meaningful contributions to FHB research, future directions will include updating the pipeline developed using 2021 images with 2022 images to improve model accuracy. Prospective research leveraging the qualities of the pipeline include the time series evaluation of FHB symptoms to study host-pathogen interactions and testing the model on additional imaging platforms, such as cell phone cameras or unoccupied aerial vehicles, to further reduce phenotyping time and hardware requirements.

In conclusion, we have demonstrated a novel FHB disease assessment pipeline capable of single spike and plot aggregate disease scoring. This system has demonstrated success across locations, years, and camera angles. Finally, the pipeline is more precise than visually in-field disease rating and is as accurate as manual image annotations for disease symptoms performed by raters. This FHB image analysis pipeline represents a breakthrough in FHB phenotyping that will be relevant and useful for studying and managing this economically important fungal disease.

## Acknowledgements

This material is based upon work supported by the U.S. Department of Agriculture, under Project numbers: 59-0206-1-198 and 59-0206-2-127. This is a cooperative project with the U.S. Wheat & Barley Scab Initiative. Any opinions, findings, conclusions, or recommendations expressed in this publication are those of the authors and do not necessarily reflect the view of the U.S. Department of Agriculture. This project was also supported by the University of Minnesota Experimental Station USDA-NIFA Hatch project MIN-22-086. Julian Cooper was supported by Bayer Crop Sciences as a Bayer Crop Sciences Fellow. We thank Susan Reynolds and all the staff, research scientists, postdocs, and graduate and undergraduate students with the University of Minnesota wheat and barley breeding groups and the Steffenson group for helping and supporting with field experiments.

---

**Figure S1. Diagram of Mineral phenotyping rover and 2021 and 2022 camera configurations.** a. Front, side, and top view of the Mineral Earth Sciences LLC. phenotyping rover. b. Rover camera configurations for 2021. Images used for model building were from cameras C, D, G, H, I, and J. c. Rover camera configurations for 2022. Images for model assessment were taken with camera C and D. d. Example image from 2021 taken with side camera C. e. Example image from 2022 taken with elevated top-down angled camera D.

**Figure S2. Example plot registration and sampled images for processing in 2022 StP.** A subset of 8 total images, 4 images per camera C and D, with even GPS spacing was selected per plot.

**Figure S3. The distribution of percent of FHB per spike for each rater for the large scale manual image annotation dataset.** The average FHB percent of the spikes annotated is shown in red for each rater.

**Figure S4. The distribution of plot aggregate percent of FHB each rater for the large scale manual image annotation dataset.** The average FHB percent of the spikes annotated is shown in red for each rater.

**Figure S5. Theoretical and observed disease distributions of the manual inter-rater reliability image annotation spike subset.**

**Figure S6. Output of the automatic image analysis pipeline for spike disease and gradability inferences.** a. The distribution of gradability probabilities before filtering at a 0.50 threshold. B. The distribution of disease at each location before filtering at a 0.50 gradability threshold. c. The distribution of gradability probabilities after filtering at a 0.50 threshold d. The distribution of FHB disease percentage at each location after filtering at 0.50 gradability threshold.

**Figure S7. The relationship between the automatic image analysis (AIA) pipeline and rater disease annotations for large scale manual image annotation (MIA) dataset of spikes.** The regression line between AIA and MIA spike FHB percentage is shown in blue.

**Figure S8. Representative examples of images from StP and Crk taken in 2022.** The original image is shown in the left column of each image pair. On the right column of each image pair, the mask of the spike segmentation model is shown in purple, the mask of the disease segmentation model is overlaid in red, and the calculated percentage of FHB detected for the spike is indicated above the image.

