## Supplemental Figure 8 for "An RGB based deep neural network for high fidelity Fusarium head blight phenotyping in wheat"

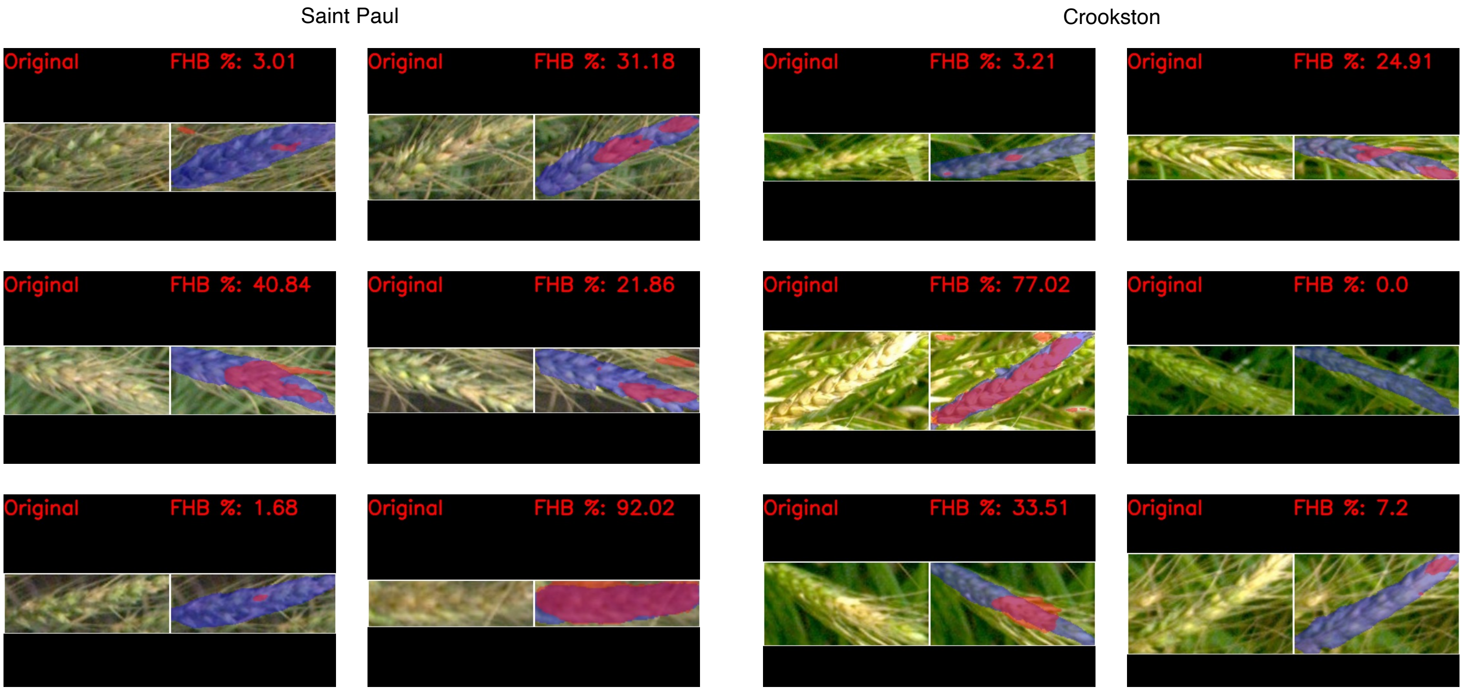


**Figure S8. Representative examples of images from StP and Crk taken in 2022.** The original image is shown in the left column of each image pair. On the right column of each image pair, the mask of the spike segmentation model is shown in purple, the mask of the disease segmentation model is overlaid in red, and the calculated percentage of FHB detected for the spike is indicated above the image.
