## Supplemental Figure 7 for "An RGB based deep neural network for high fidelity Fusarium head blight phenotyping in wheat"

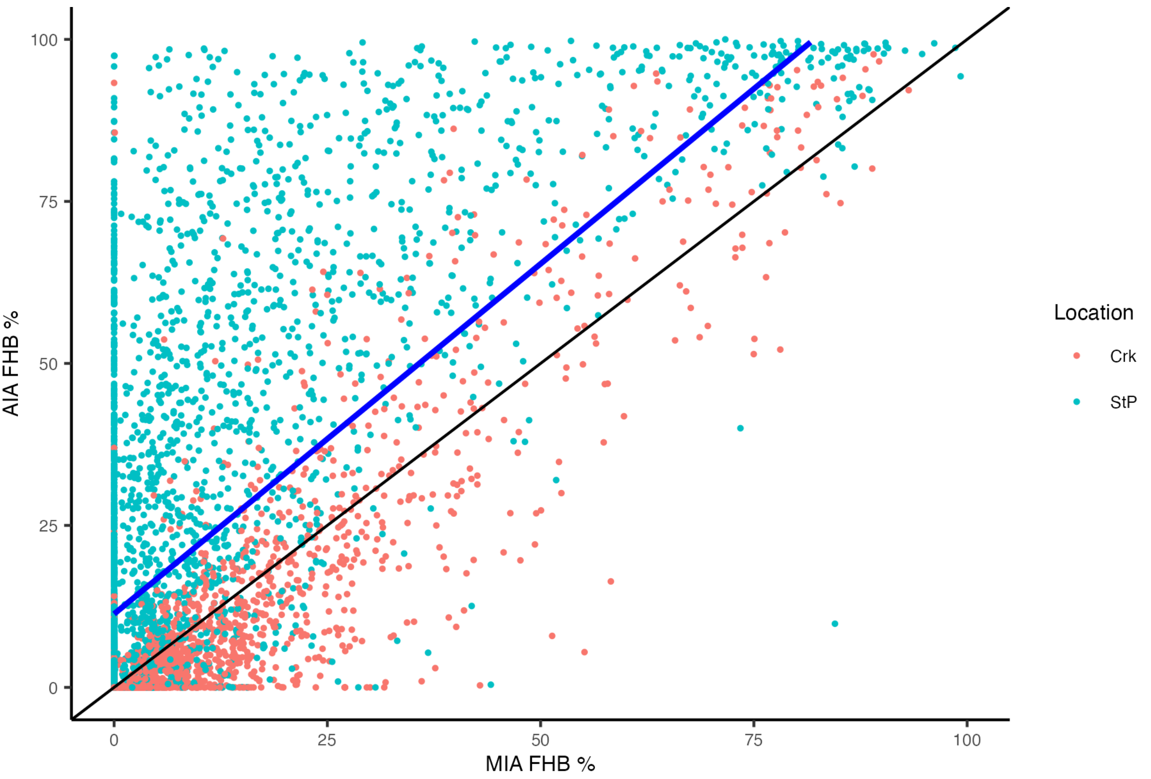


**Figure S7. The relationship between the automatic image analysis (AIA) pipeline and rater disease annotations for large scale manual image annotation (MIA) dataset of spikes.** The regression line between AIA and MIA spike FHB percentage is shown in blue.
