## Supplemental Figure 6 for "An RGB based deep neural network for high fidelity Fusarium head blight phenotyping in wheat"

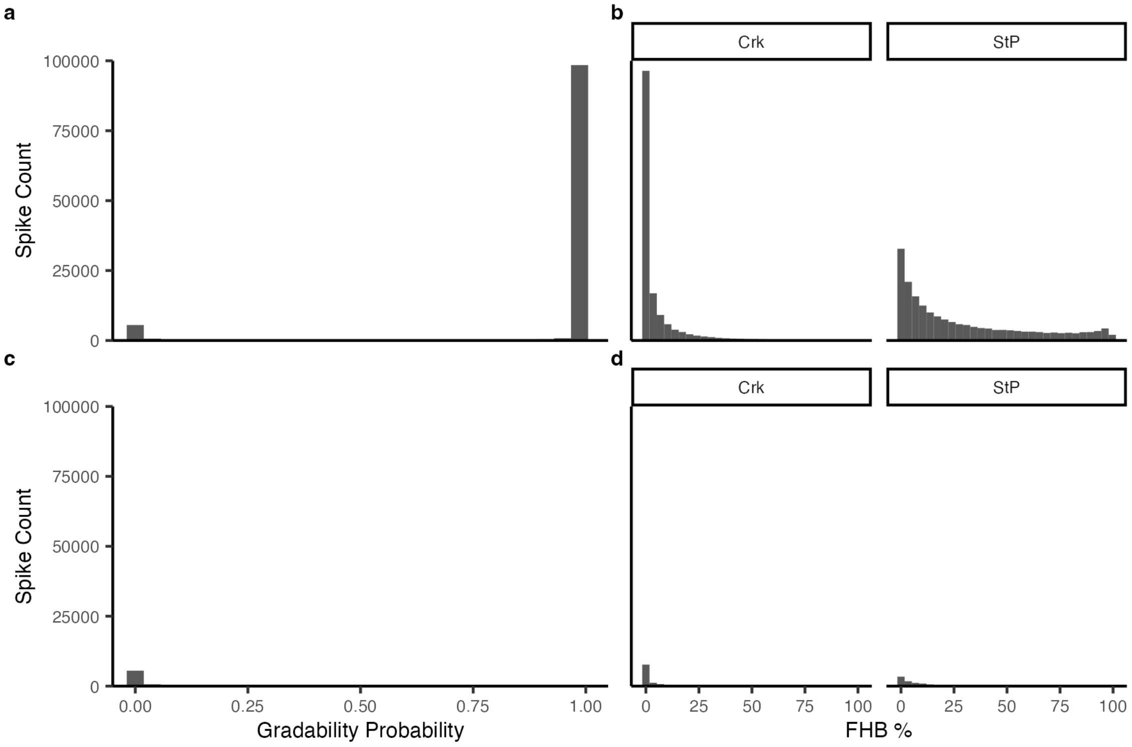


**Figure S6. Output of the automatic image analysis pipeline for spike disease and gradability inferences.** a. The distribution of gradability probabilities before filtering at a 0.50 threshold. B. The distribution of disease at each location before filtering at a 0.50 gradability threshold. c. The distribution of gradability probabilities after filtering at a 0.50 threshold d. The distribution of FHB disease percentage at each location after filtering at 0.50 gradability threshold.
