## Supplemental Figure 5 for "An RGB based deep neural network for high fidelity Fusarium head blight phenotyping in wheat"

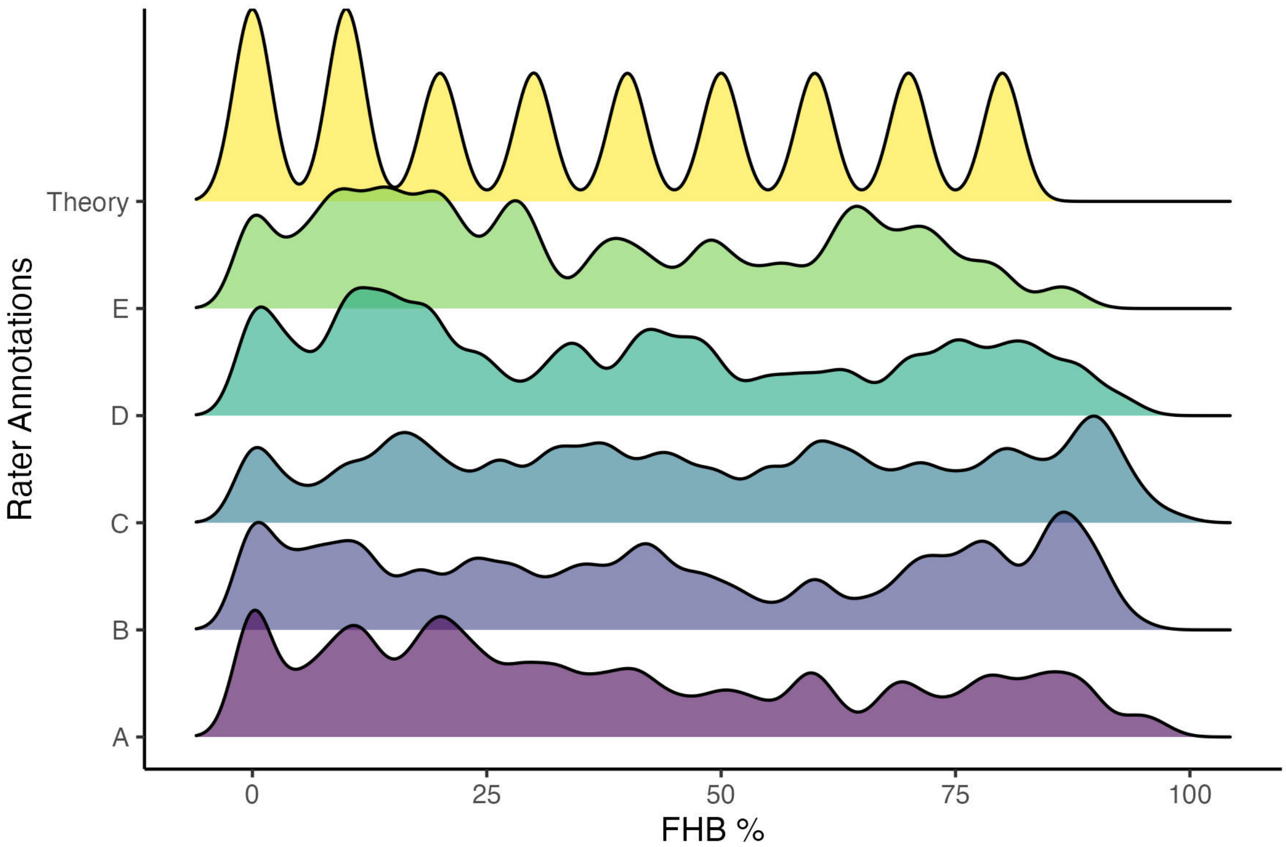


**Figure S5. Theoretical and observed disease distributions of the manual inter-rater reliability image annotation spike subset.**
