## Supplemental Figure 4 for "An RGB based deep neural network for high fidelity Fusarium head blight phenotyping in wheat"

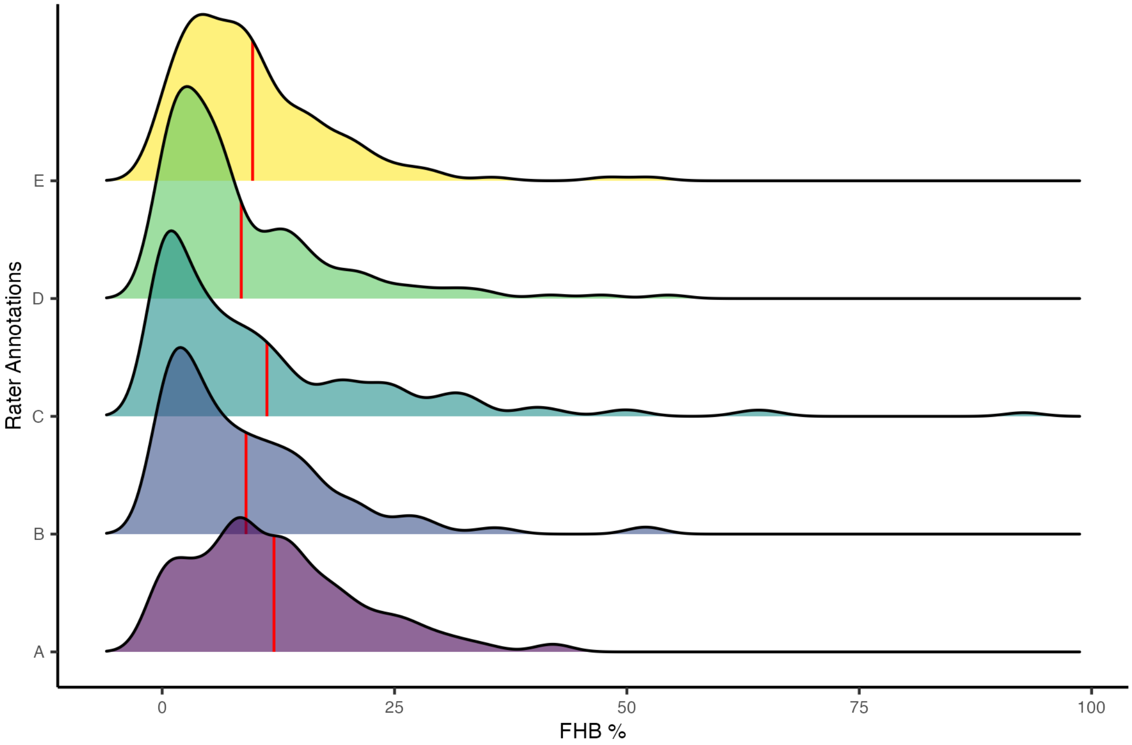


**Figure S4. The distribution of plot aggregate percent of FHB each rater for the large scale manual image annotation dataset.** The average FHB percent of the spikes annotated is shown in red for each rater.
