## Supplemental Figure 3 for "An RGB based deep neural network for high fidelity Fusarium head blight phenotyping in wheat"

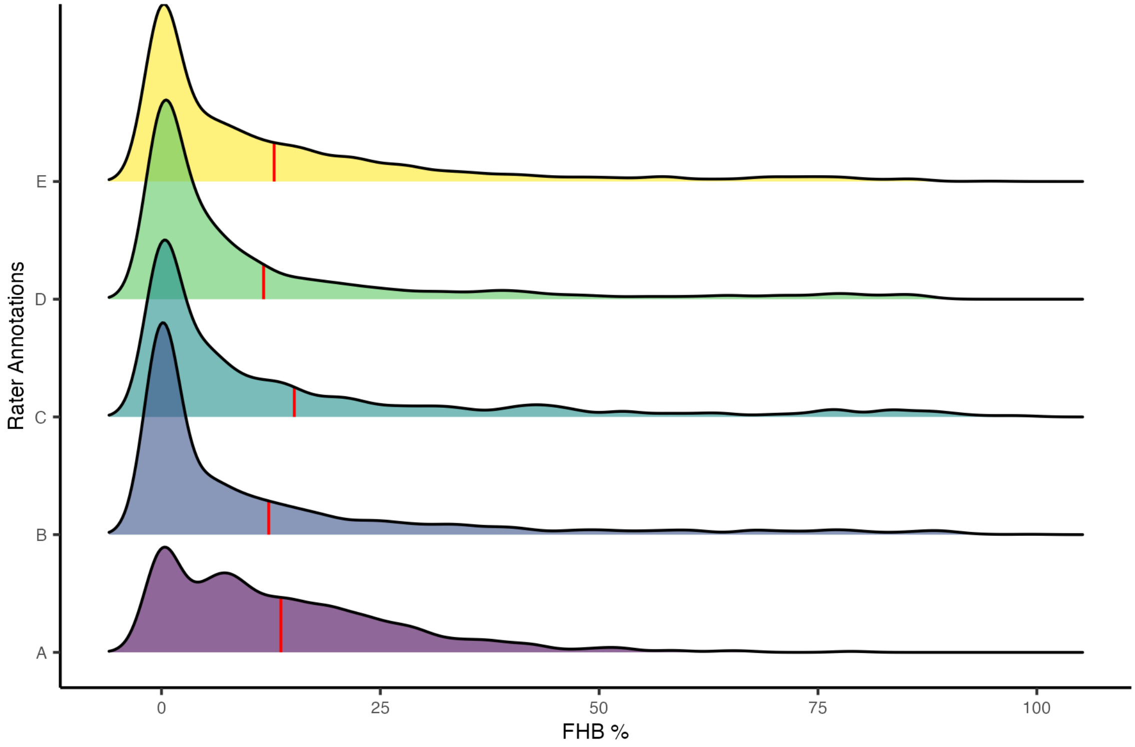


**Figure S3. The distribution of percent of FHB per spike for each rater for the large scale manual image annotation dataset.** The average FHB percent of the spikes annotated is shown in red for each rater.
