## Supplemental Figure 2 for "An RGB based deep neural network for high fidelity Fusarium head blight phenotyping in wheat"

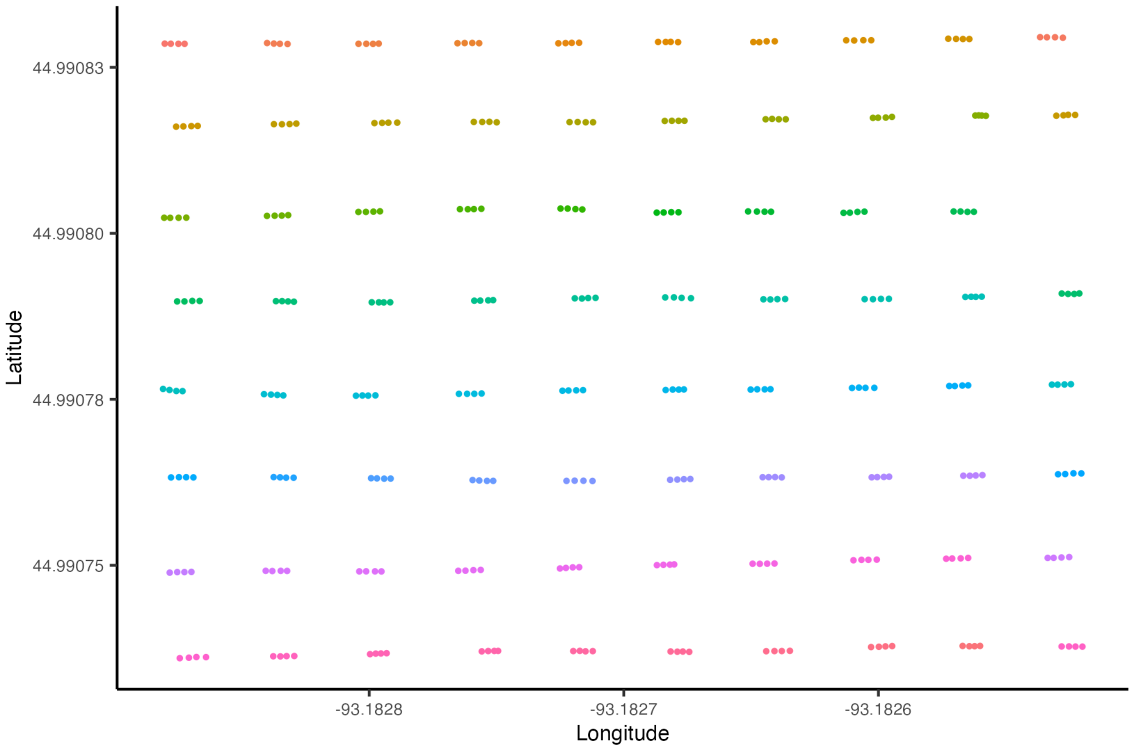


**Figure S2. Example plot registration and sampled images for processing in 2022 StP.** A subset of 8 total images, 4 images per camera C and D, with even GPS spacing was selected per plot.
