## Supplemental Figure 1 for "An RGB based deep neural network for high fidelity Fusarium head blight phenotyping in wheat"

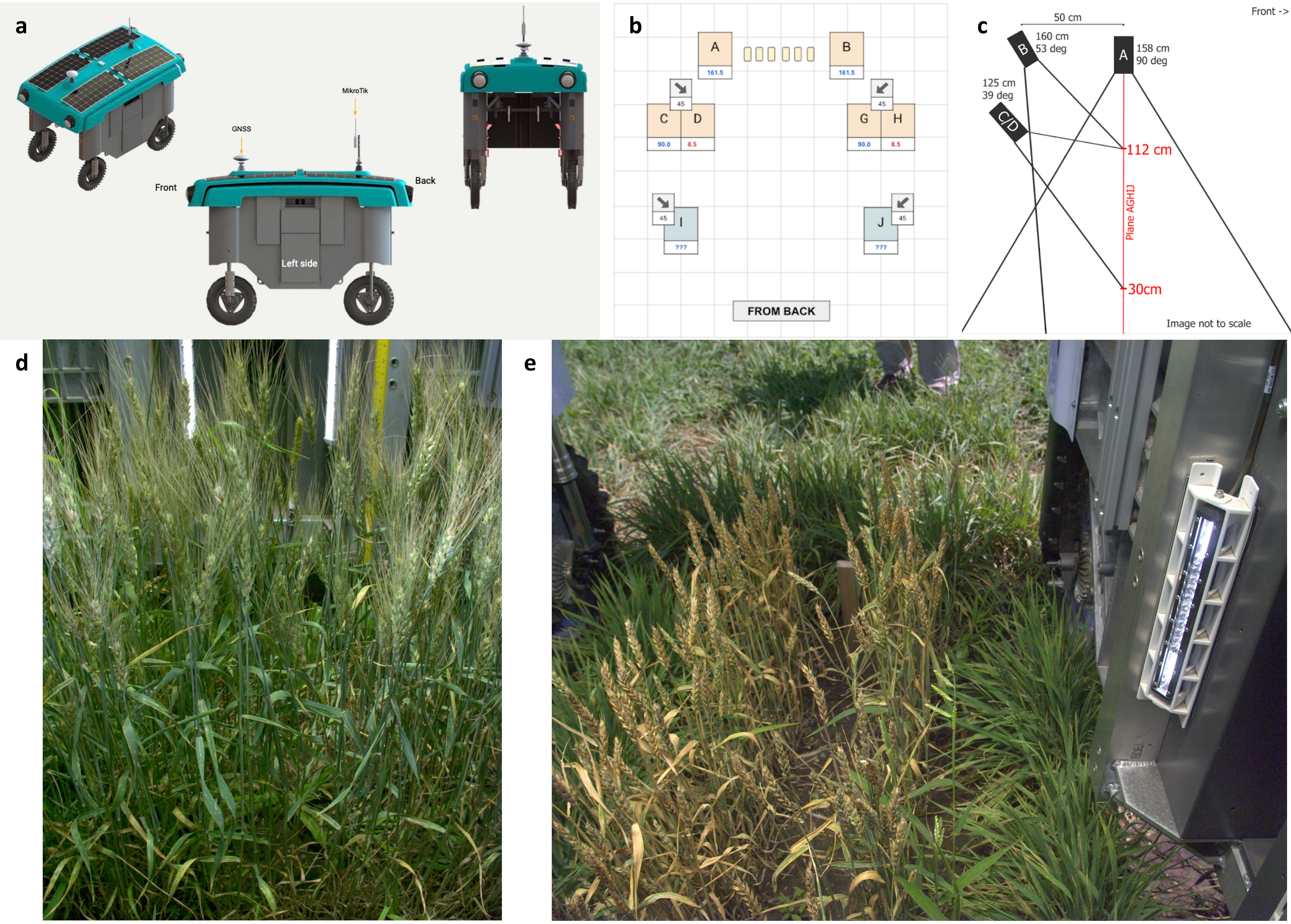


**Figure S1. Diagram of Mineral phenotyping rover and 2021 and 2022 camera configurations.** a. Front, side, and top view of the Mineral Earth Sciences LLC. phenotyping rover. b. Rover camera configurations for 2021. Images used for model building were from cameras C, D, G, H, I, and J. c. Rover camera configurations for 2022. Images for model assessment were taken with camera C and D. d. Example image from 2021 taken with side camera C. e. Example image from 2022 taken with elevated top-down angled camera D.
